# An introduction to predictive distribution modelling for conservation to encourage novel perspectives

**DOI:** 10.1101/2021.05.26.445867

**Authors:** M.P. MacPherson, K.R. Burgio, M.G. DeSaix, B.G. Freeman, J. Herbert, R. Herman, V. Jirinec, J. Shonfield, D.L. Slager, C.B. van Rees, J.E. Jankowski

**Author notes:** M.G. DeSaix (https://orcid.org/0000-0002-5721-0311). B.G. Freeman (https://orcid.org/0000-0001-6131-6832). J.E. Jankowski (https://orcid.org/0000-0003-3273-1388). J. Herbert (https://orcid.org/0000-0002-2912-2226). R. Herman (https://orcid.org/0000-0002-5914-6907). V. Jirinec (https://orcid.org/0000-0001-9856-9681). J. Shonfield (https://orcid.org/0000-0002-7159-4016). C.B. van Rees (https://orcid.org/0000-0003-0558-3674).

## Abstract

*An introduction to predictive distribution modelling for conservation to encourage novel perspectives.* The rapid pace and potentially irreversible consequences of global change create an urgent need to predict the spatial responses of biota for conservation to better inform the prioritization and management of terrestrial habitats and prevent future extinctions. Here, we provide an accessible entry point to the field to guide near-future work building predictive species distribution models (SDMs) by synthesizing a technical framework for the proactive conservation of avian biodiversity. Our framework offers a useful approach to navigate the challenges surrounding the large spatio-temporal resolution of datasets and datasets that favor hypothesis testing at broad spatio-temporal scales and coarse resolutions, which can affect our ability to assess the validity of current predicted distributions. We explain how to improve the accuracy of predictive models by determining the extent to which: 1) dispersal limitation impacts the rate of range shifts, 2) taxa are rare at their range limits, and 3) land use and climate change interact. Finally, we offer approaches to filling knowledge gaps by creatively leveraging existing methods and data sources.

**RESUMEN:** *Una introducción a la modelización predictiva de la distribución para la conservación con el fin de fomentar nuevas perspectivas*. El rápido ritmo y las consecuencias potencialmente irreversibles del cambio global crean una necesidad urgente de predecir las respuestas espaciales de la biota para la conservación, con el fin de informar mejor la priorización y gestión de los hábitats terrestres y prevenir futuras extinciones. Aquí proporcionamos un punto de entrada accesible al campo para guiar el trabajo del futuro próximo en la construcción de modelos predictivos de distribución de especies (SDM), sintetizando un marco técnico para la conservación proactiva de la biodiversidad aviar. Nuestro marco ofrece un enfoque útil para navegar por los retos que rodean a la gran resolución espacio-temporal de los conjuntos de datos y a los conjuntos de datos que favorecen la comprobación de hipótesis a escalas espacio-temporales amplias y resoluciones gruesas, lo que puede afectar a nuestra capacidad para evaluar la validez de las distribuciones predichas actuales. Explicamos cómo mejorar la precisión de los modelos predictivos determinando hasta qué punto 1) la limitación de la dispersión influye en el ritmo de los cambios de área de distribución, 2) los taxones son raros en los límites de su área de distribución, y 3) el uso del suelo y el cambio climático interactúan. Por último, proponemos enfoques para colmar las lagunas de conocimiento aprovechando de forma creativa los métodos y fuentes de datos existentes.

## INTRODUCTION

Anthropogenic pressures on climate and land cover lead to altered ecosystems and species distributions (Parmesan and Yohe, 2003). This creates an urgent need for understanding species spatial responses to global change and ensuring conservation of suitable habitat that supports population persistence and conserves biodiversity (De Frenne et al., 2021; Roberts et al., 2019). Species distribution models link species’ occurrence with ecological explanatory variables and can be used to predict range dynamics for proactive conservation measures (van Rees et al., 2022; Zurell et al., 2016); however, the accuracy of current predictive species distribution models where there is a lot of occurrence data is still limited by the variation in temporal and spatial resolution of datasets used to build them (König et al., 2021). This is because the capacity of species to change their geographic ranges is limited by both large-scale factors (i.e., range-wide drivers like temperature and rainfall that define macrohabitat) and small-scale factors (i.e., individual-specific biotic interactions and resource availability that affect microhabitat use). This issue is exacerbated for data-poor species, such as rare, cryptic and consequently, many tropical species. The limited availability of high-resolution datasets capable of capturing fine-scale dynamics thus decreases our predictive capacity for many such taxa.

A multitude of factors can ultimately influence how species, populations, and individuals respond to changing climate regimes driven by global change (reviewed in Peterson et al., 2019). Individuals may lack flexibility in tracking relevant environmental cues to adjust the timing of life history events or to otherwise accommodate novel ecological circumstances. Alternatively, even when species are flexible in timing life history events, they may be unable to modulate behavior in response to environmental anomalies that affect recruitment. For example, advancing egg-laying dates to match earlier spring warming temperatures may have negative consequences for nestling survival when the chances of cold snaps are decoupled from warming trends (Sauser et al., 2021; Shipley et al., 2020). Predictions for how animal distributions could be altered (shifted, contracted or expanded) with climate change may be species-specific (Hallman and Robinson, 2020; Radosavljevic and Anderson, 2014), but research into how particular axes of climate change affect species with different life history strategies, foraging guilds or habitat affinities provides promising insights that may apply to a wider number of unstudied species (changing rainfall regimes -Brawn et al., 2017; e.g., rearrangements of ecosystems - Huntley et al., 2008; advances in spring phenology and breeding success - Sander et al., 2020). Finally, the potential for range shifts is ultimately limited by whether potential habitat is available, accessible, and whether population density is high enough at the appropriate range edge to facilitate a shift (Block and Levine, 2021; Stiels et al., 2021).

A major gap in the current understanding of how organisms are likely to respond to global change remains in the effect of land use change and its interaction with climate change. Dispersal limitation is a critical component of distribution model theory, but is difficult to estimate in practice (Sousa et al., 2021). Species with high site fidelity (Merkle et al., 2022) or inflexible migratory routes (Stanley et al., 2012) may not be able to adopt prospecting behaviors (searching for new sites; Cooper and Marra, 2020) to accommodate dispersal to appropriate habitat following change. In the Northern Hemisphere, avian migration routes and resource tracking are well established (Faaborg et al., 2010; Thorup et al., 2017), but individuals still may not be able to correctly time migratory movements to keep up with the pace of climate change, as evidenced in declining populations that now experience mismatches in the timing of annual life history events and peaks in food availability (Møller et al., 2008; Sander et al., 2020). On the other hand, multiple Nearctic-Neotropical migrants have advanced their spring migration and breeding behavior in response to warming spring temperatures (Dunn and Møller, 2019; Pecl et al., 2017; Shipley et al., 2020), offering support to the idea of “evolutionary rescue”, where existing variation within annual cycles may yield previously unknown adaptive potential (Helm et al., 2019). Recent research on the interaction between land use and climate change points to the need to improve our understanding of interacting mechanisms underpinning risks to population persistence under climate and habitat stressors (Schulte to Bühne et al., 2021).

Here, we provide an overview of predictive bird distribution modeling with the aim to inform future work focused on how dispersal limitation, biotic factors, and abiotic factors affect distributional shifts in birds and other taxa driven by global change. Birds are excellent indicators of environmental change, and they have the most widespread and in-depth databases on distributions (Morrison, 1986; Riddell et al., 2021), making them excellent focal taxa for predictive conservation modeling. Citizen science initiatives and long-term research programs have yielded an abundance of predictive distribution literature, and insights from avian literature should be generalizable to other taxa because birds display a wide variety of life history strategies (S̜ekercioğlu et al., 2019).

We synthesize theory, methods, and data sources widely used to predict bird distributions under global change to highlight best practices and opportunities for innovation and refinement. Understanding where species occur and why some areas are occupied but not others is essential for developing effective conservation plans. Predictive distribution modeling is important to build upon our knowledge of distributions based on surveyed areas and to identify potential areas for protection or management. Modeling wide ranging, unevenly distributed, cryptic, or difficult to survey species all present problems for understanding factors driving occupancy. Here, we bring together the issues that often complicate predictive distribution modeling to offer a solutions-oriented approach to habitat conservation under global change.

## ECOLOGICAL & EVOLUTIONARY THEORY AS A BACKBONE TO APPROACHING PREDICTIVE SDMs

The first step in our synthesis is to illuminate the importance of a taxon’s ecology and evolution in determining their spatial response to global change. Dramatic changes in climate over the remainder of this century are expected (Figure 1), with many species likely unable to survive in all of the areas they presently occupy (Ceballos et al., 2015; Sax et al., 2013). Species typically respond to climate change in one of three ways: 1) by going extinct, 2) persisting via shifts in geographic range, or 3) adapting in place via evolutionary change. Evidence from the fossil record suggests all three responses are common (Graham et al., 1996; Smith et al., 1995). The capacity to adapt *in situ* in response to climate change has been relatively understudied (Jirinec et al., 2021; Parmesan and Matthews, 2005). A meta-analysis of observed phenological and morphological adaptations to climate change concluded that morphological trait adaptation, while variable among taxa, will not likely work on a sufficient scale to mitigate the worst effects of ongoing climate change, whereas phenological adaptation (e.g., timing of reproduction or migration) could mitigate such effects, albeit imperfectly (Radchuk et al., 2019). Whether species with relatively shorter generation times should be better able to survive by evolving in place remains understudied. Advancing such research is hampered by the difficulty of conducting experiments on large numbers of species at sufficient time scales needed to develop a comprehensive framework (Holt et al., 2005; Pierson et al., 2015). Variability among taxa and the short time scales in which we have been making observations together indicate that selection pressure has yet to be strong enough to produce measurable changes, though it appears that species shift their phenology faster where the rate of climate change is higher (Poloczanska et al., 2013). Precipitation, and to a lesser extent temperature, are the primary drivers of adaptive responses to climate change (Caruso et al., 2017; Jirinec et al., 2021; Siepielski et al., 2017).

**Figure 1.**
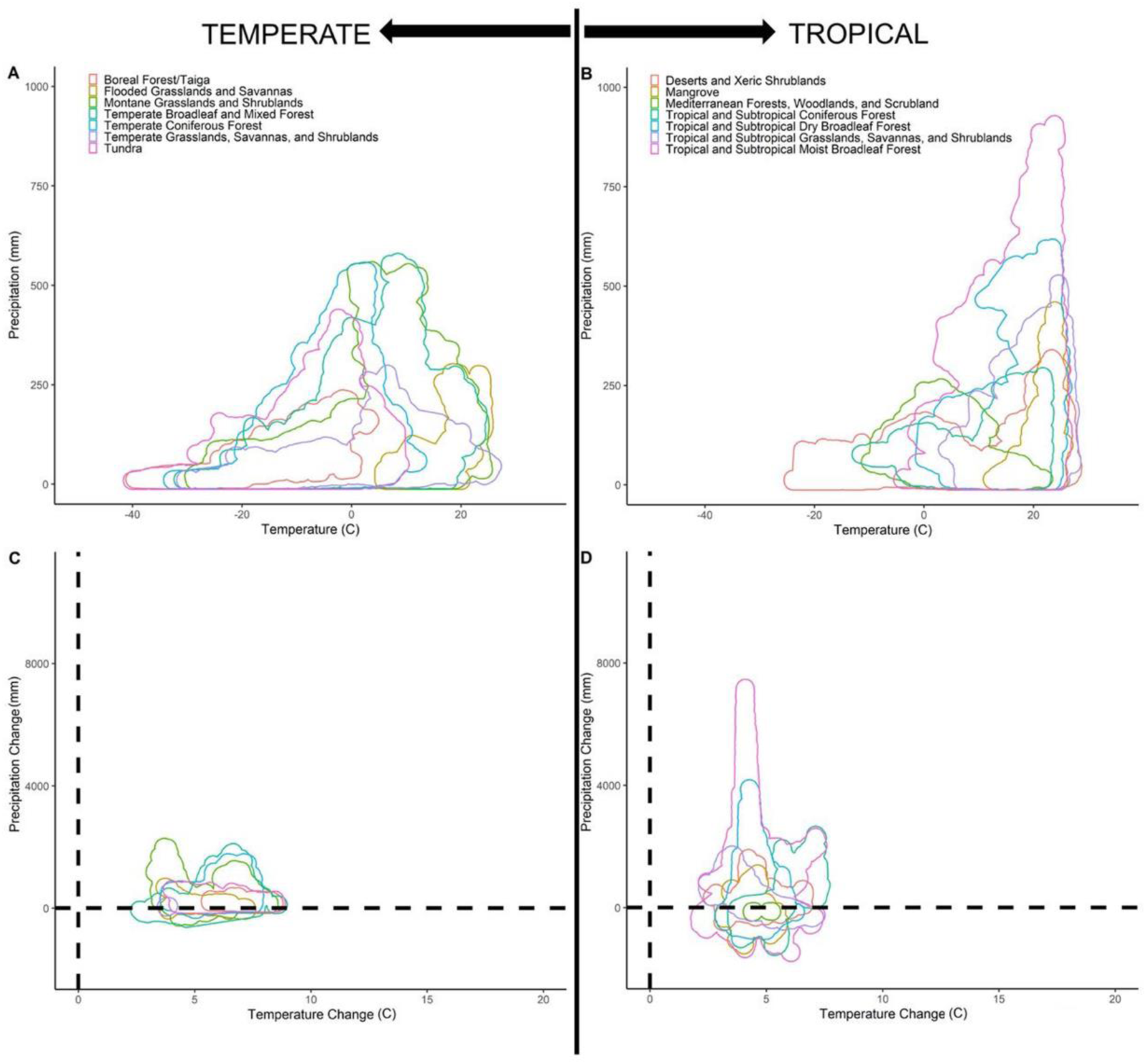
Current climate and predicted change in climate of global terrestrial biomes. Climographs of (A) temperate and (B) tropical and subtropical global terrestrial biomes. Projected changes in temperature and precipitation regimes for (C) temperate and (D) tropical and subtropical global terrestrial biomes. While some areas are predicted to have increased or decreased precipitation in the future, no areas are expected to have lower or stable temperatures. Temperature increases are predicted to be higher in temperate biomes, while changes in precipitation are greater for tropical biomes. Climographs built from The World Wildlife Fund terrestrial ecoregions (https://www.worldwildlife.org/publications/terrestrial-ecoregions-of-the-world) and current climate (2021-2040) from CanESM5 middle of the road scenario (SSP3-7.0) accessed from Worldclim (https://www.worldclim.org/data/cmip6/cmip6climate.html). The projected change is modeled as the difference between current climate (2021-2040, described above) and future climate using the worst-case scenario (SSP4-6.0; 2081-2100) from CanESM5 global climate model data (Meinshausen et al., 2020). Species’ presence, and thus biodiversity, is highly influenced by biome type, which shifts from one form to another according to temperature and rainfall regimes (Whittaker 1975). Projected changes in temperature and precipitation may cause global terrestrial biomes to shift in space, but spatial gains and losses by different biomes over time may not occur in synchrony with habitat needs to maintain biodiversity (Dorazio et al., 2015). R code for producing this figure can be found at [https://github.com/mmacphe/Global_Change_Biomes].

Relevant ecological covariates inform one of two main modeling categories to predict where a taxon could occur: 1) species distribution models (hereafter SDMs) that comprise presence records and abiotic data, and 2) ecological niche models (hereafter ENMs) that explicitly estimate the accessible environment (Soberón and Peterson, 2005). These approaches are meant to have high spatial accuracy but are not intended to inform on cause-and-effect species-habitat relationships (Merow et al., 2013). Whether using SDMs or ENMs, these models can be based on different niche perspectives, which present unique frameworks for estimating drivers of occupancy across a species’ range. Here, we list simplified descriptions of three niche concepts to provide a basic introductory framework for understanding the domain of predictive distribution modeling as all are used in modern predictive modeling (see *An Overview of Analytical Methods* section for more information). Models based on the Grinnellian niche concept focus on abiotic drivers of site occupancy (Elton, 1927; Soberón, 2007; Wisz et al., 2013) and SDMs are mainly built under this framework (Figure 1; Saupe et al., 2012). Coarse-scale variables describing ecosystem characteristics are often the most relevant for predicting shifting distributions across large spatial extents. In comparison, the Eltonian niche concept includes estimates of biotic interactions and resource-consumer dynamics that are only quantifiable at local scales (Figure 1; Elton, 1927; Soberón, 2007). Finally, the Hutchinsonian niche concept reflects the functional role of a species, which is often estimated using functional traits and habitat requirements based on functional traits (Rosado et al., 2016), and projects the probability of site occupancy beyond study areas, including estimates of dispersal probabilities in ENMs (Figure 2). Because a species’ ability to move beyond their current realized niche depends on the relative importance of abiotic factors, biotic interactions, and dispersal probability, which can operate at different spatial scales (Jankowski et al., 2013), knowledge of these categories is important for making sound inferences from whatever data and analytical approach are used.

**Figure 2.**
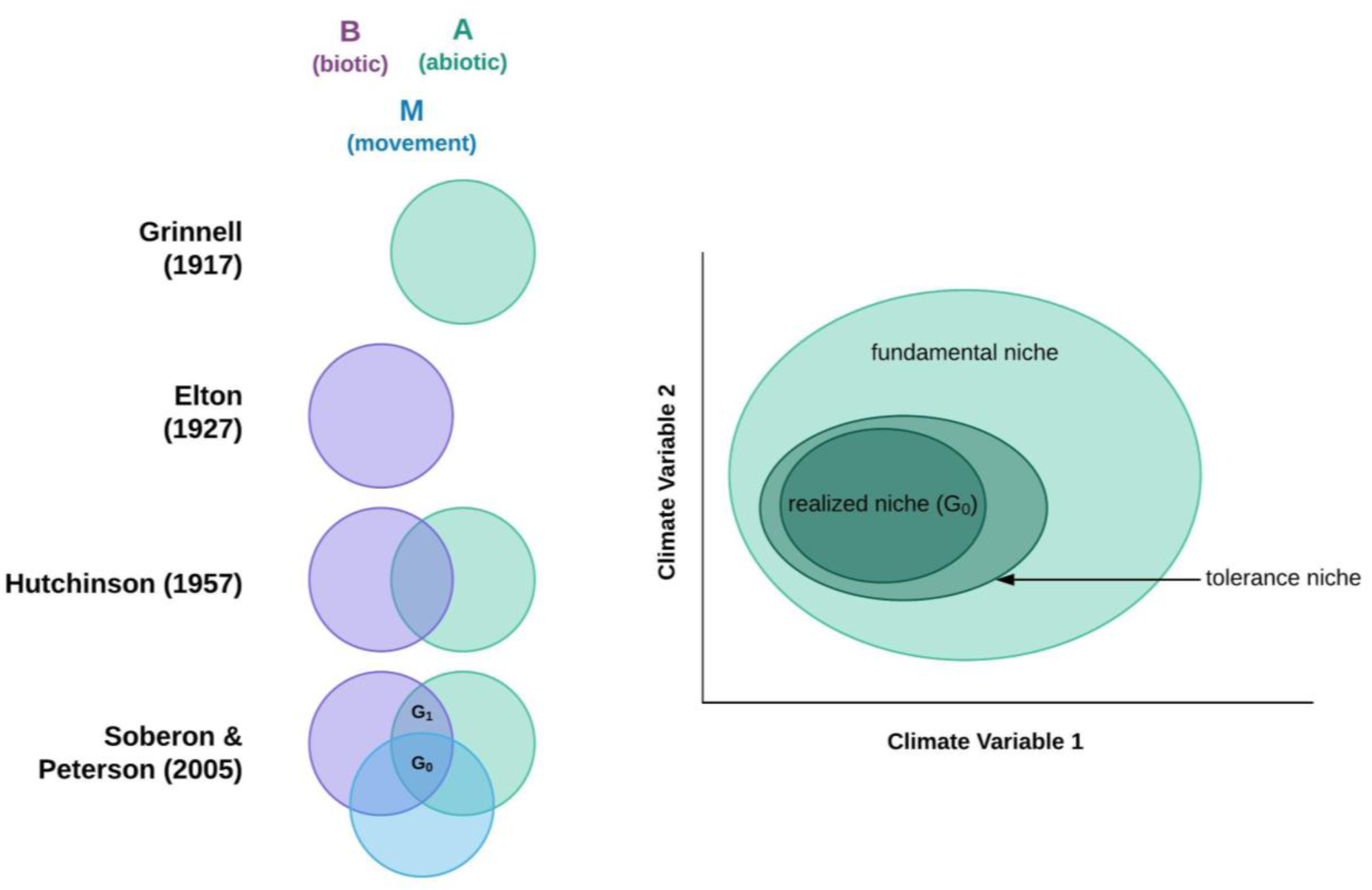
The “BAM” diagram. The relationship between “B” - biotic, “A” - abiotic, and “M” - movement (also thought of as “access” and defined by dispersal ability) with respect to the authors of various niche theories (left). The location of realized, tolerance, and fundamental niche along the intersection of two hypothetical climate variables (right). Sax et al. (2013) present an update to niche theory that considers three components of a species niche: 1) the “realized niche”, i.e., conditions within the native range, 2) the “fundamental niche” - conditions, in which a species could thrive if it were introduced there, and 3) the “tolerance niche” - conditions, in which individuals could survive, but likely unable to maintain populations over the long-term. The fundamental niche is the set of appropriate abiotic conditions, and the realized niche is a smaller area where both abiotic and biotic environments are suitable. Theoretically not all of the realized niche could be accessible and so the potential range is yet smaller because it reflects only the area that is suitable and also accessible to the species. Occurrence is not expected outside of the colored areas. The occupied area is represented by G0, and G1 reflects appropriate conditions that have not or cannot be accessed by the species (as described by Saupe et al., 2012).

While ENM tools can be used to predict spatially beyond the current range of a taxon, they are only able to predict the probability of site occupancy within the range of abiotic conditions measured within current range boundaries. Thus, assessing occurrence or abundance along abiotic gradients is important for predicting future range shifts under global change, as this relates directly to the response of species or populations to the environment. The gradual shifts in temperature and rainfall regimes can be expected to align with changes in species responses, reflecting the shifts in optimal ambient conditions and resulting in altered patterns of probability of occurrence or abundance. However, ENM tools are limited where climate change introduces novel conditions (e.g., Williams et al., 2007), and more mechanistic (rather than correlative) analytical approaches may provide more accurate predictions.

Demographic information is also useful to incorporate for more accurate predictions because spatial ecology theory dictates that species respond to continuous environmental gradients through gradual changes in abundance, as individuals experience shifting conditions toward or away from environmental optima over space (Austin, 2007, 2005). Populations may also show thresholds in abiotic tolerances or in response to biotic factors that change their response shape (e.g., creating asymmetric or skewed response functions; e.g., Oksanen and Minchin, 2002). For example, the asymmetric abiotic stress limitation (AASL) hypothesis predicts a steeper decrease in a species’ probability of occurrence toward the more stressful end of a species’ distribution, which has been supported in several vascular plant species (Dvorský et al., 2017). Other threshold-type effects can result in asymmetric responses shown across multiple species in communities (e.g., sharp ecotone boundaries, appearance of dominant predators or competitors; Jankowski et al., 2013). Identifying response shapes is of practical interest to understand the impact of abiotic and biotic factors on occupancy. The evaluation of responses along abiotic gradients are becoming more common but often rely on high spatial resolution survey data (Bani et al., 2019; Burner et al., 2019; e.g., Maggini et al., 2011; Urli et al., 2014) that is difficult to amass. However, with more studies, comparisons across species would enable distinguishing commonalities and differences in responses to gradients and could point to key environmental variables driving patterns in community organization.

Whether populations would be able to respond (in time and space) to changing conditions also depends on heterogeneity within landscapes; this is the realm of landscape ecology land use change. Landscapes can vary naturally or due to anthropogenic alterations in ways that affect both the phenology of resources (time) or availability of habitat (space); such variability can be tracked if conditions change at a pace that populations could respond to. Examples of this include species adapted to annually ephemeral or temporally patchy habitats such as wetlands and grasslands; species accustomed to such environments may have dispersal strategies that predispose them to better track climate (e.g., *Elaenia cristata* - Ritter et al., 2021). In other cases, habitat and resources are spatially patchily distributed, and the ability of species to respond to changing conditions may be more taxon-specific for those that rely on spatiotemporal heterogeneity that is naturally patchy. Examples of this include habitat specialists in naturally patchy habitats such as Amazonian white sand habitats (e.g., Rufous-crowned Elaenia (*Elaenia ruficeps*) - Ritter et al., 2021).

Alternatively, animals may avoid the problem of range shifting altogether by buffering themselves from acutely unfavorable conditions using microclimate refugia. One example of this is through behavioral thermoregulation. Ambient temperature is almost never constant in terrestrial environments—it varies by time, and by habitat, often at very small scales (Scheffers et al., 2017, 2014). Biologists have long understood that mobile animals exploit thermal heterogeneity to maintain optimal body temperature (Angilletta Jr and Angilletta, 2009; Angilletta et al., 2009; Cowles and Bogert, 1944; Porter et al., 1973; Stevenson, 1985). For cold-adapted species in a warming world, this can be achieved by shifting activity times to cooler periods of the year and day, and by moving to microhabitats with cooler microclimates. This capacity to compensate for unfavorable ambient temperature by behavioral thermoregulation, known as the “Bogert Effect” (Huey et al., 2003), can at least partly mitigate the harmful effects of climate warming (Huey et al., 2012). However, behavioral thermoregulation is contingent on access to cooler areas and periods; animals already occurring in the most buffered environments have limited options for escape when conditions change. Furthermore, individuals could experience additional pressure from biotic interactions with species moving into their habitat to thermoregulate (Huey et al., 2012).

Whether taxa will be able to respond to global change is ultimately a taxon-specific question driven by the constraints of dispersal limitation; one of three fundamental aspects of distribution theory (Figure 2). Understanding barriers to distributions caused by dispersal limitation has been of great interest, for example, in bird species unable to cross areas of unsuitable habitat, and has been studied both experimentally (e.g., Moore et al., 2008; Naka et al., 2022) and theoretically (Ribas et al., 2018, e.g., 2012). Selection pressures driving morphology, such as migratory compared to sedentary life histories, may play a role in physically limiting dispersal distance (Capurucho et al., 2020; Claramunt et al., 2012; MacPherson et al., 2022; Sheard et al., 2020), even in birds. Life history strategies also dictate the demographic trends in dispersal; for example, in many species it is the inexperienced young that disperse into new areas away from the territories of their parents (e.g., Florida Scrub Jay (*Aphelocoma coerulescens*) - Suh et al., 2020). Dispersal limitation is thus a complex and difficult-to-quantify aspect of predictive distribution modeling that is nevertheless fundamental for predicting the probability of range shifts with global change (reviewed in Zurell, 2017). Coupling studies of distribution modeling with quantifiable measures of dispersal capacity is an important next step toward making more informed predictions of species responses to changing environments (Sousa et al., 2021; Travis et al., 2013; Urban et al., 2013).

Ultimately, extinction risk (and conversely, population viability) centers on declining population and small population paradigms; population ecology. Population declines are driven by deterministic (demographic, abiotic, and biotic) factors that cause reduction to small numbers (e.g., the Allee effect), and also by stochastic factors, which dominate the extinction dynamics of small populations (Caughley, 1994; Morris and Doak, 2002; Smith et al., 2021). The abiotic shifts in conditions due to climate change and the resulting shifts in species distributions play a role in both types of population dynamics. Shifting distributions can cause temporary or permanent reductions in habitat area (Thomas et al., 2006; van Rees and Reed, 2018), lowering carrying capacity and driving population declines at local or regional scales, or reducing connectivity and creating isolated subpopulations that are more susceptible to small-population dynamics (Anderson et al., 2009). The degree to which changing distributions result in commonly observed conservation impacts depends on a diversity of species traits including habitat preferences and associations, phenotypic plasticity, and current range and population size (Jiguet et al., 2007; Visser, 2008). Notably, research based solely on changes in distributions tends to predict widespread extinctions due to the aforementioned mechanisms, but plasticity, dispersal, and adaptation appear to have been key for many species’ resilience to past and current climate change (Moritz and Agudo, 2013). For example, the Galapagos finches are a well-known example of phenotypic plasticity in bill morphology in response to environmental change (Grant and Grant, 2020; Grant and Weiner, 2017; Weiner, 1994).

Conservation action that occurs at small spatial scales requires deep natural history knowledge for decision-making. In this section we described the domain of how ecological theory informs spatial responses to land use and climate change. When designing research projects that aim to predict future distributions under the umbrella of conservation ecology, we urge modelers to deeply consider the ecology of focal taxa by referring to this section again in the future, and the references cited within it. **Box 1** summarizes essential broad considerations for predictive distribution modeling by the novice modeler.

### Box 1. Essential Broad Considerations for Predictive Distribution Modeling by Novice Modelers.

Here we summarize this section by highlighting common pitfalls observed by the authors while acting as reviewers in the peer-review process.

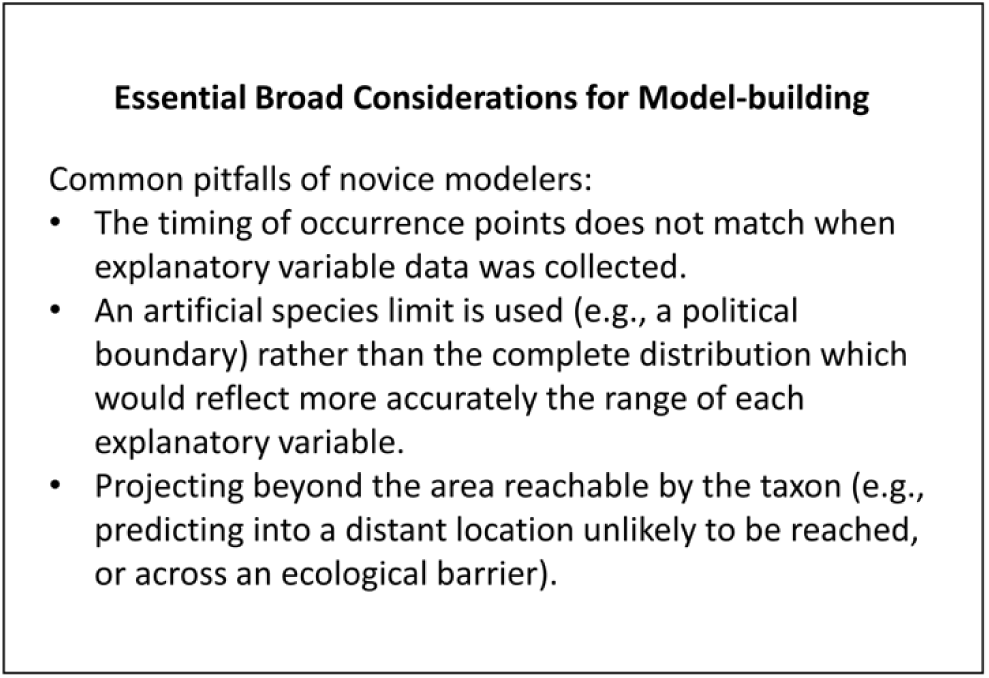

## A TECHNICAL OVERVIEW OF OCCURRENCE DATA FOR PREDICTIVE DISTRIBUTION MODELING

Occurrence data through time is the foundation for applied predictive distribution modeling. Here, we present technical aspects of different sources of avian occurrence data to inform which questions can be answered with different data sets. To assess whether a change in distribution has occurred or will occur, researchers require historical records to establish a baseline for comparison, followed by surveys at one or more subsequent time points. Although birds are among the best studied class of animals, early baseline inventories are still rare, and often consist of field notes (Reznick et al., 1994), which present challenges particularly in reconstructing past abundances (Shaffer et al., 1998). The aim of general collecting for birds by museums is to document with vouchered specimens the complete taxonomic representation of a location, thus supplying presence and absence information. A caveat to this is that complete information is limited to small areas of intense investigation, thus limiting the widespread application of data gathered by museums in presence/absence modelling (e.g., Loiselle et al., 2003). Although general collecting for birds is no longer the norm at some museums (Ferguson, 2020; for reasons, please see Remsen, 1995), museum records remain the best sources of historical occurrence data, though they primarily supply presence or presence/absence data only and not abundance data (Shaffer et al., 1998).

A major challenge for predictive models is rarity or sparseness of point occurrence data. Even with many potential sources of occurrence data, knowledge of bird distributions still varies greatly by species, habitat, and geography. For example, rare, nocturnal, and/or secretive species and those in remote areas are difficult to detect and model accurately (MacKenzie et al., 2005; Stralberg et al., 2015). Rarity is a common characteristic of species in diverse tropical communities and remains a major challenge for constructing species distribution and environmental niche models for tropical species (e.g., Marini et al., 2010).

In contrast to sparse information from the past, modern ornithologists benefit from technological advances that offer unprecedented information on bird occurrence including from devices that track individuals beyond specific field sites (satellite imagery - Borowicz et al., 2018; and sophisticated automated biodiversity data collection - Kitzes and Schricker, 2019; for birds within appropriate size ranges - McKinnon and Love, 2018; Sullivan et al., 2014, online citizen science efforts - 2009). They improve our understanding of avian ecology by identifying previously unknown species-habitat relationships (Jirinec et al., 2016), migratory routes (Jahn et al., 2021, 2016; Stanley et al., 2015, 2012), foraging areas and behaviors (including accessing microhabitat refuges - Wolfson et al., 2020), and wintering distributions (Renfrew et al., 2013). These advances facilitate discovery of species responses to global change by revealing locations outside of direct observation. Aside from emerging technologies, simpler monitoring approaches implemented in the past at large scales have amassed rich long-term legacy datasets (e.g., breeding bird surveys, regional atlases, and the 5 million bird eggs housed at natural history museums - Marini et al., 2020), and band recoveries across the globe continue to supply information on bird population trends and distributions.

Decisions about which type of occurrence data is best for predictive modeling hinges on the trade-off between addressing processes affecting the core distribution versus a more holistic understanding including unique responses at range boundaries, during different life history activities, or between different sex and age classes. Passive acoustic monitoring and long-term studies remain promising approaches for capturing relative abundance data to estimate population dynamics with respect to global change (Pérez-Granados et al., 2019; Pérez-Granados and Schuchmann, 2020; Sugai et al., 2020). Trait databases are gaining momentum as sources of functional diversity to test hypotheses under the Hutchinsonian niche concept (Gallagher et al., 2020; Leclerc et al., 2020; Matuoka et al., 2020; Tobias et al., 2022). Exploring how functional roles of species change across abiotic gradients can dramatically improve our understanding of abiotic tolerances to more accurately predict range dynamics. For example, bill size has been shown to be important for thermoregulation in birds (e.g., Danner and Greenberg, 2015) and a study of functional trait structure along a tropical elevational gradient in Malaysian Borneo linked larger bills in low elevation communities with thermal tolerance (Boyce et al., 2019). Current shortfalls in avian occurrence data (summarized by Lees et al., 2020) require an integrated approach to predicting bird distributions with global change. Although most efforts to collect occurrence data are from small spatial areas (e.g., from individual field research sites) and coarse resolutions (Table 1), this data contributes meaningfully to our understanding of species-habitat relationships that can be applied to predictive modelling.

**Table 1.** Geospatial context of common occurrence data sources for birds and explanatory variables. Most occurrence data and explanatory variable data layers are of coarse spatial and temporal resolution, making them useful for testing hypotheses under the Grinnellian class of niche. Occurrence data sources with fine spatial and temporal resolution are not as common and require higher investment for acquiring location data (i.e., GPS tags) or physically tracking individual birds (i.e., radio-tracking) but could be important for testing hypotheses under the Eltonian class of niche.

| Geospatial data qualities | Examples of occurrence data sources | Examples of explanatory variables |
| --- | --- | --- |
| Large spatial area, coarse spatial and/or temporal resolution | •Remote sensing assets (satellite imagery, aerial imagery etc.)<br>•Automated radiotelemetry<br>•Geolocators<br>•Citizen Science: eBird, European Bird Census Council, European Breeding Bird Atlas, South African Bird Atlas Project<br>•Stable hydrogen isotopes<br>•Genetic data<br>•Natural History collections | •Long-term averages in stable climates (e.g., WorldClim)<br>•Land cover (e.g., MODIS, GlobCover, SPOT)<br>•Phenology and vegetation indices (e.g., MODIS, Landsat)<br>•Climate record (e.g., NCDC, ClimateNA, ClimateSA)<br>•Atmospheric weather reanalyses (e.g., ERA – European Centre for Medium-Range Weather Forecasts Re-Analysis, GPM – Global Precipitation Measurement, TRMM – Tropical Rainfall Measuring Mission, NCEP – National Centers for Environmental Prediction) |
| Small spatial area, coarse spatial and/or temporal resolution | •Citizen science (e.g., Breeding Bird Surveys, Christmas Bird Counts)<br>•Natural history collections<br>•Field notes |  |
|  | •Point count surveys<br>•Passive acoustic monitoring<br>•Long-term datasets<br>•Bird banding data |  |
| Large spatial area, fine spatial and/or temporal resolution | •GPS tags | •Digital elevation models (e.g., ASTER)<br>•Vegetation indices (e.g., SPOT) |
| Small spatial area, fine spatial and/or temporal resolution | •VHF radio-tags | •Personal weather data loggers<br>•Weather station data |

## SELECTING APPROPRIATE EXPLANATORY VARIABLES

Predicting distributions relies on accurate assessments of life history requirements. In this section, we synthesize the spatial and temporal resolution of explanatory variables commonly used to match distributions with habitat and resource needs. Producing sound predictive models requires ecologically relevant, suitable proxies for resource or habitat needs (e.g., Burns et al., 2020; Randin et al., 2020). To scale up to entire distributions, there must be an underlying knowledge of a species’ life history that effectively links the lives of individuals to the explanatory variable (or proxy variable) associated with site occupancy.

Predicting distributions influenced by changing food webs requires a breadth in knowledge of life history information to predict the probability of species’ ranges shifting individually or with their entire biotic community. This task is complicated by the potentially differing needs for each life history stage or demographic, which is often the case for birds (e.g., habitat requirements may differ between fledglings and adults). While main considerations are the spatial and temporal resolutions of explanatory variables, biotic interactions can also play a critical role (see Fig. 1) and may need to be considered on a taxon-specific basis.

### CONSIDERING THE SPATIAL RESOLUTION OF EXPLANATORY VARIABLES

Selecting the correct spatial resolution of explanatory variables to use with occurrence data is key to producing accurate assessments of species-habitat relationships. First, matching the spatial resolution of explanatory variables with our understanding of the scale at which individuals make occupancy decisions (e.g., whether they are micro-or macrohabitat based) is important for making proper predictions of where organisms could move in the future. Second, matching spatial resolution between explanatory variables and occurrence data prevents pseudoreplication. Specifically, occurrence data may be pseudoreplicated if many samples occur in the same pixel of explanatory data. Data spatial resolution also affects the possibility of model overfitting (i.e., when the importance of a narrow set of conditions is overestimated because occurrence data come from a limited portion of conditions actually experienced by the taxon).

Remotely sensed spatial products (e.g., NDVI and BIOCLIM variables) can be proxies for features of the environment being used by individuals at macrohabitat scales that require occurrence data from a wider distribution. Remote sensing systems with finer spatial resolution, for example light detection and ranging (LiDAR), have been used to predict occupancy by light-demanding versus shade-tolerant plant species (Wüest et al., 2020), and direct measurements of structural features of habitat (e.g., vertical distribution of forest canopy elements and foliage density) used to predict occupancy (e.g., Goetz et al., 2010). Remotely sensed variables are thought to reflect a contingency of resources assuming the data product conveys mutual information on a shared spatiotemporal scale with the amount of food resources (Riotte-Lambert & Matthiopoulos, 2020). There are some satisfying interpretations of the ecological relevance of these remotely sensed products (Bridge et al., 2016 (EVI); Renfrew et al., 2013 (NDVI); see Tøttrup et al., 2012 (drought)), but the typical purpose of using these large scale variables is for spatial accuracy and not to infer causal relationships.

However, explanatory variables that reflect some ecological relevancy is important for building accurate predictions of how taxa may respond to global change. Some remotely sensed abiotic products do reflect a direct habitat resource or type. Habitat characteristics (i.e., land cover) are most often identified with remotely sensed data, which depending on the sensor, can have a variety of resolutions. Medium-high spatial resolutions (≤30m, e.g., Landsat, Sentinel) are useful to pair with more precise occurrence data (e.g., GPS tags, point counts; Shirley et al., 2013), whereas low spatial resolution (e.g., MODIS) can be useful for occurrence data from geolocators or weather radar (Heim et al., 2020). The temporal resolution of remotely sensed data is a limitation, especially when modeling the dynamic nature of migratory animals in seasonal environments (MacPherson et al., 2018a; Roslin et al., 2021).

### CONSIDERING THE TEMPORAL RESOLUTION OF EXPLANATORY VARIABLES

The temporal predictability of resources is important for predicting future distributions because annual life history strategies are dependent upon correctly timing life cycle events with required resources. For example, it is widely held that birds breeding at high latitudes must correctly time the hatching of young with annual insect emergence to maximize fledging success, and early arrival to breeding grounds may enhance reproductive success (Alerstam, 2011; Kokko et al., 2006; Nilsson et al., 2013; Smith & Moore, 2005). Predictive distribution models are expected to be the most accurate when species-habitat correlations are assessed using occurrence and environmental data gathered from the same time-period. However, the temporal resolution of occurrence data is much finer than that of environmental data; this can create a mismatch that limits the questions that can be answered using current approaches. The habitat requirements for fulfilling life history needs may vary depending upon the life history event (i.e., mate acquisition, rearing young, molting, migrating) and the species life history strategy. Life history strategies vary in birds from those that have evolved to rely on consistent resource availability (e.g., in the case of dietary or habitat specialists that cannot live outside of narrow environmental conditions), to periodic resource abundance (e.g., seasonal, annual, inter-annual weather patterns that drive migratory or irruptive population movements), or irregular resource availability (e.g., in nomadic taxa), and it is important to match locality data in time to environmental data for accurate species-habitat assessment.

The temporal resolutions for widely used climate data varies, with trade-offs between temporal and spatial resolution in addition to the state of products limiting the types of distribution questions that they can inform (Table 1). We expand on this using the examples of temperature and rainfall explanatory variables in Appendix 1 because these are the two most widely used and generally important factors in species distribution modeling (Bradie and Leung, 2017).

To build robust predictive models, it is necessary to match the spatial and temporal resolution of the occurrence data with that of the habitat characteristics. The temporal resolution of remotely sensed data is a limitation, especially when modeling the dynamic nature of migratory birds where dynamic seasonal habitat changes drive short-term habitat quality (e.g., seasonally flooded mudflat habitat for migrating shorebirds; Twedt, 2013). Google Earth Engine (a javascript-based platform where one can write their own code to integrate many satellite products and independently calculate indices such as NDVI) is a flexible tool that can facilitate identifying the correct temporal resolution of land cover data to robustly test explanatory power of land cover variables. Examples of data sets in Google Earth Engine include vegetation indices (Landsat, ∼2 weeks temporal resolution), products for creating land cover classifications (SPOT,∼1 month), and static land cover datasets (e.g., GlobCover, Cropland Data Layers, National Land Cover Database; Table 1).

### CONSIDERING BIOTIC FACTORS

Modeling future range shifts in response to global change rarely considers biotic factors. This omission is the main reason why an assumption of SDMs - that the species is at equilibrium with their environment - is rarely, if ever, met (Pearson and Dawson, 2003). If this assumption were supported, the realized and fundamental niche would completely overlap as long as the species could access all available suitable niche space (Figure 1). Consideration of biotic factors has the potential to significantly enrich the field of predictive distribution modeling with more accurate forecasts, and here we synthesize leading research contributing to this end at both the intraspecific and interspecific level.

It is thought that biotic interactions vary along abiotic gradients such that they can either enhance or reduce predicted ranges (Louthan et al., 2015). However, because capturing this information requires both high spatial and temporal accuracy across all abiotic scenarios within the realized niche, there are few examples to draw from. One promising approach to including biotic interactions in predictive spatial modeling is the development of causal models that estimate the influence of interspecific competition using co-occurrence data (e.g., Staniczenko et al., 2017). The scope and strength of biotic factors may be correlated with abiotic pressures (Louthan et al., 2015), and may differ depending upon which part of the range is being considered. For example, much work has been done in the bird literature to improve our understanding of species-habitat relationships through the study of hybrid zones, which typically occupy only one part of a species range (eg., Taylor et al., 2014). Quantification of biotic factors, like the ones described here, requires targeted species-specific research to test hypotheses within the Eltonian and Hutchinsonian niche concepts. This research is mainly done within small spatial extents and without consideration of temporal or spatial resolution (e.g., co-occurrence records from different time periods or datasets with different spatial resolutions - Atauchi et al., 2018; Palacio and Girini, 2018; see also König et al., 2021), but some research has been done at large spatial extents and coarse resolutions (e.g., Acorn Woodpecker (*Melanerpes formicivorus*) occurrence with Colombian Oaks in the Northern Andes - Freeman and Mason, 2015).

Intraspecific competition can drive density-dependent range shifts in migratory species such that interannual selection of locations are dependent on group size and food availability (Corre et al., 2020). Dispersal limitation can contribute to inadequacies in recruitment to new or population sink sites, limiting population recovery or range shifts despite available habitat (Palma et al., 2020; Zurell et al., 2016). The ability to update behaviors when circumstances change (behavioral flexibility) may be key for driving range shifts or expansions under global change (Blaisdell et al., 2021). Further, bird species capable of behavioral innovation (a.k.a. plasticity) often have lower risk of extinction, as they are better able to adapt to changing ecosystems and habitat destruction (Ducatez et al., 2020; Reed et al., 1999). However, metapopulation dynamics can affect the introgression of adaptive traits such that maladaptive traits restrict range shifts under global change (e.g., Garcia-R and Matzke, 2021; Lavretsky et al., 2020). Each of these examples are implicitly expected to vary across abiotic gradients, highlighting the importance of assessing the probability of site occupancy beyond the core of a species distribution.

Perhaps more difficult is estimating the strength of interspecific biotic factors in shaping distributions because this requires a breadth of natural history knowledge beyond that of the species in question. However, interspecific biotic factors are thought to have strong influences on the probability of site occupancy. Species may be excluded through predator/prey dynamics where patches of habitat occupied by a species’ predator may be less likely to be occupied by the species being studied (Léandri-Breton and Bêty, 2020). Interspecific competition can also exclude species and is often assessed using co-occurrence data (Elsen et al., 2017; Freeman et al., 2016; Jankowski et al., 2010). Commensalisms, where one organism benefits from the presence of another organism and the other unaffected, can also facilitate site occupancy (Aitken and Martin, 2007). Including data on the presence of predators, competitors, and commensalisms in SDMs can be a useful addition to improving models (Jankowski et al., 2013). Alternatively, the spatial overlap with other taxa may have little influence if the focal species exhibits behavioral avoidance (e.g., via interference competition - Jaeger, 1970; or predator-prey dynamics - Lukas et al., 2021) or the spatio-temporal scope of data are not aligned.

The utility of SDMs to guide predictions of distributions under global change will certainly be limited by the spatio-temporal resolution of environmental variables. Even when researchers are mindful when selecting environmental data suited to answer questions about their occurrence data (i.e., using ecologically relevant environmental and climate data across biologically relevant space and timescales), a challenge remains in combining and contrasting the Grinnellian niche concept (e.g., climate variables) against other niche concepts that require finer temporal and spatial data (e.g., density of interacting species). This problem can further complicate predictive models if site occupancy is dependent on time-lags, as is the case in some migratory species (Bridge et al., 2016), and is explored further in the following section.

## AN OVERVIEW OF ANALYTICAL METHODS

Predictive models of species distributions can be divided into correlative versus mechanistic approaches (Figure 3). Correlative models elucidate relationships between species occurrence and spatially explicit explanatory variables and can be viewed as hypothesis-generating tools. Data to parameterize correlative models are readily available (e.g., correlating occurrence data with land-cover or weather data as described above), making them the dominant approach to predictive distribution modeling. Mechanistic models (a.k.a. causal models) are hypothesis-testing tools that incorporate physiological tolerance to predict where species will be capable of persisting within their physiological limits (Guisan and Zimmermann, 2000); data to support these approaches is more limited. Here, we summarize popular SDM tools with important information to help novice modelers identify which tool might be appropriate given their data and question. We provide examples that students can refer to for accessing more detailed methodologies.

**Figure 3.**
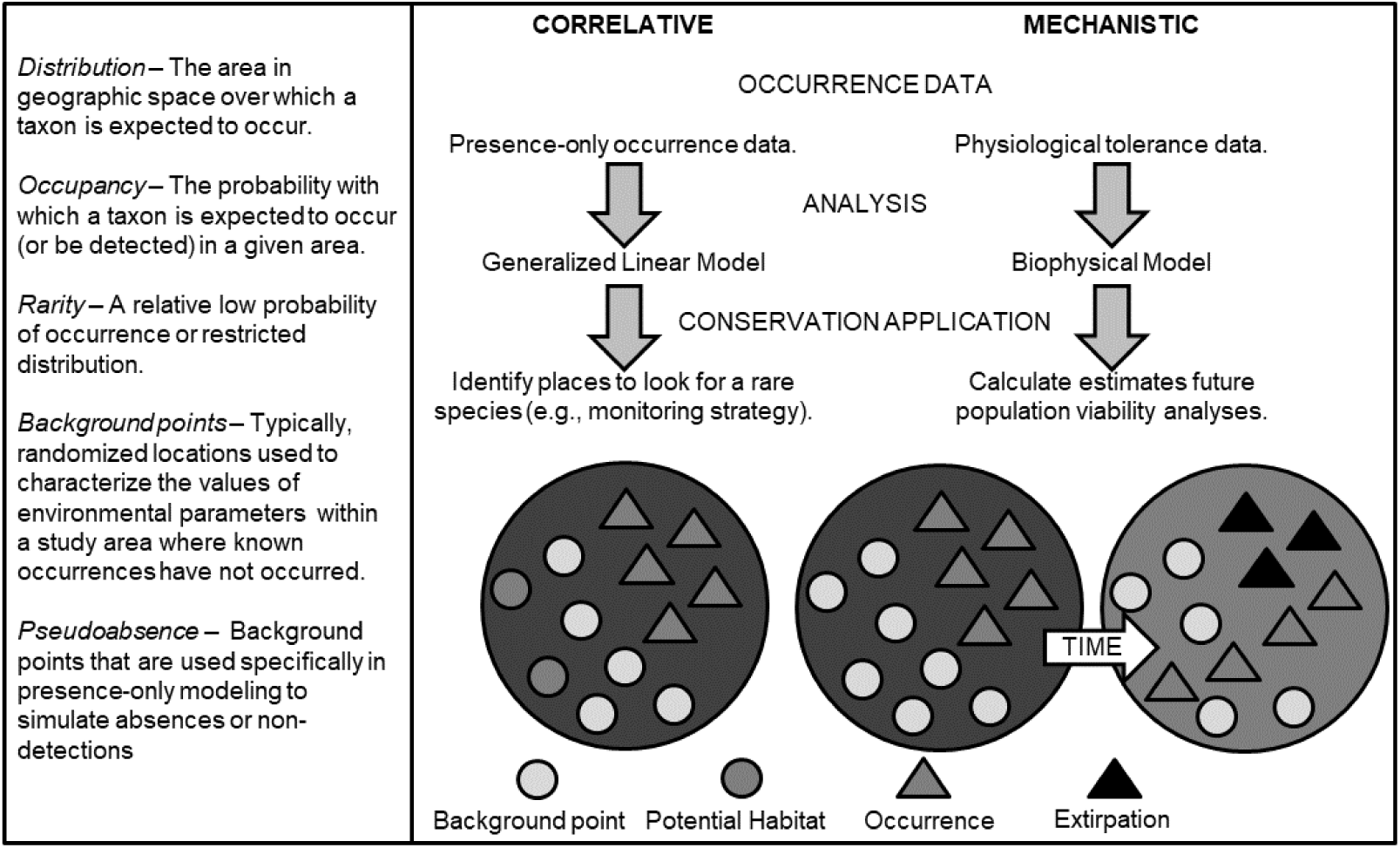
Analytical processes for species distribution models. Left: important terms used in describing distribution model data. Right: Parallel workflows for example correlative versus mechanistic predictive modelling.

Range estimates from correlative SDMs are valuable, helping us better understand species and their sensitivities to global change. MAXENT is one of the most popular SDM approaches, widely used for its ability to use presence-only datasets where true absence or non-detection data are typically hard to find (Elith and Leathwick, 2007; Merow et al., 2013). MAXENT has been useful for modeling distributions of rare or cryptic species due to its apparent high performance with small or incomplete datasets (Aubry et al., 2017; Pearson et al., 2007; Phillips et al., 2006; Wisz et al., 2008; but see also Marini et al., 2009). Boosted regression trees (BRTs) are one of many machine learning techniques well-suited to modeling complex ecological data because they can handle different types of predictor variables, fit complex nonlinear relationships, accommodate missing data and automatically handle interaction effects between predictors (Elith et al., 2008; Graham et al., 2008). BRTs have been used to predict bird distributions using occurrence data for land birds (Veloz et al., 2015), waterfowl (Barker et al., 2014), shorebirds (Dalgarno et al., 2017), seabirds (Oppel et al., 2012), and owls (Domahidi et al., 2019). Hierarchical Bayesian models can also integrate various occurrence data types (e.g., presence-absence, presence-only, count data) to create reliable spatio-temporal distribution models (Hefley and Hooten, 2015). Using continuous spatial predictor variables such as land cover, the abundance and distribution of species can be predicted over a continuous latent surface (Chakraborty et al., 2010), which has been useful with waterfowl aerial survey data (Herbert et al., 2021, 2018). Generalized linear models (GLMs) and generalized additive models (GAMs) can be employed to understand distributions across multiple environmental conditions and also conceptualize patterns in distributions of abundances across environmental gradients (Augustin et al., 1996; Smith & Edwards, 2021). GAMs are used to model nonlinear relationships (e.g., Maggini et al., 2011). One drawback to GAMs, however, is that evaluation of response shapes (e.g., optima, skewness) is done through visual inspection rather than statistically. Huisman-Olff-Fresco (HOF) models are used to predict occupancy along abiotic gradients and are particularly useful in evaluating alternative shapes of species responses to gradients, for example, if changes in abundance do not exhibit the widely assumed symmetric Gaussian functions, but instead show asymmetric, threshold-like changes in occupancy over space (Huisman et al., 1993). HOF response curves can be used to highlight distinct distribution patterns such as species replacements, allowing inference of potential biotic interactions that are otherwise difficult to measure (Jansen and Oksanen, 2013). Species climate response surfaces (CRSs) model the probability of occurrence using bioclimatic variables (Huntley et al., 1995). Given that explanatory variables in correlative SDMs cannot be used to infer causation, the CRS approach can accommodate interactions between abiotic variables to predict distributions under future climate scenarios (Huntley et al., 2006). The choice of predictive SDM technique and quality of SDM product are based predominantly on the type of occurrence data available and secondarily on spatiotemporal matching of explanatory variables.

Mechanistic approaches to SDMs predict areas where physico-chemical processes meet life history needs (Kearney, 2006; Kearney & Porter, 2009). For example, a biophysical model for the endangered Night Parrot (*Pezoporus occidentalis*) in Australia shows how air temperature increases of 3 °C would lead to lethal levels of dehydration (Kearney et al., 2016). Bayesian networks (BNs) are another type of mechanistic model that go beyond species-habitat correlations by also considering processes that influence occupancy across space and time (i.e., site access and selection - Jones, 2001). Originally employed to model human judgement (Pearl, 1985), BNs have only recently been adapted for predictive SDMs (MacPherson et al., 2018b; Staniczenko et al., 2017). Mechanistic models are informed by causal relationships based on empirical research or expert knowledge. The latter are referred to as belief networks (Drew and Collazo, 2014; MacPherson et al., 2018b). Mechanistic models enable the identification of the most important variables driving distribution patterns through mapping the fundamental niche, which is helpful to inform conservation and management decisions when circumstances change.

Ensemble modeling strategies, in which the predictions of multiple approaches are combined or used simultaneously, are often suggested to better encompass the range of uncertainty in prediction (Araújo and New, 2007). This method accounts for the fact that model choice is often the greatest source of quantifiable uncertainty in species distribution modeling and reduces sources of bias from the use of a single algorithm (Dormann et al., 2008; Jarnevich et al., 2017). Implementing ensembles of five or more of the algorithms and approaches described above has been greatly facilitated by software packages like BIOMOD (Thuiller et al., 2009) and VisTrails SAHM (Morisette et al., 2013), which bring methods together into a single analytical environment. Ensembles can be used to assess a range of potential projected outcomes, forming a “bounding box” or “consensus” across algorithm predictions (Araújo and New, 2007). Analysis with model ensembles typically involves scaling and averaging model outputs and, often weighting these by some measurement of model performance such as “area under the curve” (AUC) score.

The ongoing “big data revolution” in many fields including ornithology (La Sorte et al., 2018) has increased use of artificial intelligence for data handling and analysis (Xia et al., 2020), especially in correlative distribution modeling. Machine learning algorithms like MAXENT (Merow et al., 2013), random forests (Mi et al., 2017), neural networks (e.g., Manel et al., 1999), deep learning (Benkendorf and Hawkins, 2020) and boosted regression trees (Elith et al., 2006) are now commonly applied SDMs, and are valued for prediction and interpolation. They are excellent tools for insight into potential future shifts under global change (Elith, 2017).

Conservation plans rely on SDMs for future species distributions, despite the small, but growing, body of literature on ways to incorporate climate change into conservation planning (Alexander et al., 2017; Hole et al., 2011; Jones & Cheung, 2015; Loyola et al., 2013; Nakao et al., 2013; Terribile et al., 2012; Willis et al., 2009). The predictive power of SDMs for informing conservation decisions remains limited by a lack of tool capability to calculate confidence intervals around predictions (Marini et al., 2010), a lack of validation to reduce uncertainties and the need for agreed-upon standards for guiding model building (Araújo et al., 2019, 2005). Using correlative predictive models is likely to overestimate range shifts and extinction risk due to the violations of common assumptions in correlative distribution modeling. For example, SDMs tend to violate the assumption that the individual species is currently at equilibrium within its environment (Early and Sax, 2014; Sax et al., 2013) and do not take into account species interactions (Pearson and Dawson, 2003). Despite these criticisms, SDMs remain the “best” widely accessible approach currently in use for identifying potential future habitat-area (Tingley et al., 2014).

## DISCUSSION

To invite novel perspectives into the field of predictive distribution modelling, we reviewed the theory underpinning how global change shapes range dynamics and outlined the technical details of data sources and analytic approaches needed for creating sound tests of ecological theory to improve predictive models. Predicting where species will live in an uncertain future is a central goal of modern ecology. This effort has recast the basic research question of what limits distribution of species – why a species lives *here* but not *there* – as an applied question. Species are already on the move in response to recent decades of global change (and, depending on the region, predicted trends of reduced or increased rainfall; Figure 2), supporting long-held theory that climate limits species’ geographic ranges (Tingley et al., 2009). However, the match between species’ distributions and climatic conditions is often weak (Chapman, 2010; Suggitt et al., 2012); species vary considerably in their observed responses to recent climate change (Lehmann et al., 2020; Mamantov et al., 2021). These observations indicate that we are still in the early stages of ecological forecasting. To guide future research, we identified three core motivating questions that build from our technical introduction as the answers to these questions are particularly likely to generate important advancements in the field of predictive distribution modeling.

1. *Is dispersal a rate-limiting step in range expansions?* The proximate driver of range shifts is dispersal; ranges expand when individuals move beyond the existing range limit. Dispersal constraints are thus one obvious explanation for cases where the rate of species’ observed range shifts are failing to track with the rate of changing climate. Birds have incredible capacities for dispersal (e.g., Slager, 2020), making it tempting to underestimate the possible role for dispersal constraints in avian range shifts, including unexplored factors such as site fidelity. However, even bird species often show extreme limitations in dispersal, suggesting that longer-distance range shifts (e.g., across latitudes) may be slow, particularly in heterogeneous landscapes, or when species show high specialization to their associated habitats. In the temperate zone, even some long-distance migrants may show strong site fidelity to breeding (and wintering) territories (Winger et al., 2019), which suggests the possibility that rapid range shifts may be difficult to achieve. Lastly, dispersal may be possible but may introduce a new trade-off with other components of the annual life cycle, such as migration. For example, the ability of boreal breeding birds to take advantage of newly available forest habitat in the Arctic may be constrained by a trade-off with ever-increasing migration distances. We can improve our understanding of dispersal limitation by creatively pairing historical with contemporary datasets to identify patterns of dispersal limitation in places where global change has already altered species distributions.
2. *Are species particularly rare at their range limits?* Ecological theory assumes that species are most abundant at the center or core of their range and become progressively rarer towards their range limits (the “abundant-center hypothesis”; Brown, 1984). However, empirical patterns of abundance often fail to fit these expectations. Whether the abundant-center hypothesis generally holds or not is now a matter of debate (e.g., Dallas et al., 2020), and the answer may have consequences for rates of species’ range shifts. For example, low abundances at range edges could lead to slower rates of range shifts than if abundance distributions are more uniform across a core-to-limit transect through a species’ geographic range. This could be due to simple numeric effects – range shifts are easier when there are more individuals at the range edge to start with – or to genetic effects. That is, low abundances (and hence lower genetic diversity) could reduce local rates of adaptation at range edges, with gene flow from the abundant range center swamping any effect of local adaptation at the range edge that would facilitate range expansion when conditions change (Haldane, 1956; but see Kottler et al., 2021). Pairing occurrence data with natural history knowledge of species expanding their ranges has the potential to improve our understanding of the mechanisms that promote expansion from original range boundaries (e.g., the tolerance niche of the Asian Openbill (*Anastomus oscitans*) - Lei and Liu, 2021; and climate matching in the European Bee-eater (*Merops apiaster*) - Stiels et al., 2021). Additionally, assessing occupancy from the perspective of abiotic gradients rather than focal taxa would improve our ability to identify significant geographic barriers to range shifts with climate change.
3. *How does land use change interact with climate change?* Like most of the range shift literature, we have focused in this review on predicting species’ responses to changes in climate per se. But species are shifting (or failing to shift) their ranges in landscapes that are increasingly dominated by human activities (e.g., Fumy and Fartmann, 2021). Studies that simultaneously incorporate both land use and climate change as drivers of distributional change are few in number but hold great promise for several reasons. First, land use change is an obvious driver of species’ distributions; many species simply do not persist in landscapes with extreme levels of human control. Second, appropriate habitat for most species in human-dominated landscapes consists of habitat patches of varying sizes with varying levels of connectivity (e.g., Neilan et al., 2019). This fact elevates the role of dispersal in determining whether patches (and scaled-up to a larger geographic scale, regions) are occupied by a given species. Third, habitat change directly affects local-scale climate. Fragmented forests, for example, average substantially warmer and drier than primary forests (Kapos, 1989; Nunes et al., 2022), and “urban heat islands” alter local temperatures that are known to affect bird distributions (in migrants - Bonnet-Lebrun et al., 2020; and residents - Latimer and Zuckerberg, 2021). Hence, land use change can act as a multiplier of temperature effects on species. Here we emphasize the importance of high spatial and temporal resolution datasets for identifying mechanisms of global change that affect individual organisms or populations. In light of this, mechanistic (rather than correlative) models are likely to be better suited to identifying the interaction of land use and climate change when environmental conditions can be linked to the biological processes of individuals (or the loss of migration with increased urbanization - Bonnet-Lebrun et al., 2020; e.g., thermoregulatory behaviors of individuals in unshaded desert areas - van de Ven et al., 2019).

The consequences of climate change for species are potentially severe, with widespread predictions of extinctions (Thomas et al., 2006), yet the application of predictive models to conservation and management are still limited (but see Casazza et al., 2021). Some evidence suggests these dire predictions may be coming true. For example, warming temperatures have led to local extinctions in mountaintop communities in southeastern Peru as species shift their geographic ranges to track climate, a potential harbinger of the possible extinction of high elevation tropical species (Feeley et al., 2012; Freeman et al., 2018; Rehm and Feeley, 2016). Yet species may be more resilient than models assume; for example, genetic data indicate that many species were able to persist through dramatic climate fluctuations in the Pleistocene (Bocalini et al., 2021; Song et al., 2020; Wogan et al., 2020). Application of predictive distribution modes in conservation and management should become more widespread by including the development of tools for calculating confidence intervals (Marini et al., 2010) and validating models (i.e., comparing past to present distributions with respect to global change) for correlative models, and increasing the use of mechanistic models.

Humans have already made Earth hotter than it has been since before the Pleistocene (∼ 2 mya). This rapid change in Earth’s climate has set species on the move and led to a plethora of scientific research aimed at predicting where species will live in the coming decades as warming continues. These predictions, though frequently made, are seldom tested (Tayleur et al., 2015; but see Tingley et al., 2009; Wilson et al., 2018). Without this crucial step of model validation, it is impossible to assess the usefulness of predictive models. Here, we have presented theory and highlighted a range of data sources and analytical approaches that can be used to predict species’ geographic responses to climate change. Ever-greater computational power, combined with increasingly large datasets of species occurrence and landscape covariates, permit the use of greater complexity in models (e.g., Lurgi et al., 2015). However, more complex does not necessarily equate to better. The litmus test for any predictive model is how well it predicts reality. To this end, we advocate for an increasing focus on collecting empirical data that matches the spatio-temporal resolution of occurrence data with environmental variables, tests hypotheses beyond the Grinnellian niche concept, and directly evaluates model predictions (e.g., using mechanistic SDMs or resurveys). For example, the long-term predictions from models whose predictions are not met over the short-term are unlikely to be helpful (see Willis et al., 2009). Determining species’ resiliency will depend on accurate estimates of the fundamental niche including which attributes make some species more vulnerable than others, which abiotic gradients are the most important to consider for the promotion of population persistence, and including biotic interactions in predictive, validated models. Our review aims to support the ongoing pursuit of more meaningful predictive distribution models under land use and climate change that will be of great near-term utility.

## ACKNOWLEDGEMENTS

The authors declare that we have no sources of conflict of interest affecting the objectivity of the presented topic. This manuscript is the result of a synthesis on how ornithologists predict bird distributions under global change that was presented in a series of talks at the North American Ornithological Congress, 2020. We sincerely thank the organizers of this conference for their support throughout the COVID-19 pandemic. Thank you also to the participants of the Predicting Bird Distributions Under Global Change symposium, including William Lewis. This research was funded in part by a U.S. Geological Survey Northwest Climate Adaptation Science Center award G17AC000218 to C.B. van Rees. The manuscript was approved by the Director of the Louisiana State University Agricultural Center as manuscript number 20213-XXX-XXXX. The authors would also like to thank (in alphabetical order by last name) Anna Borne, Nicholas Mason, Quinn McCallum, Diego Ocampo Vargas, Samantha Rutledge, Subir Shakya, Philip Stouffer, David Vander Pluym, and Brenna Wells who provided important discussion of the topics presented in this manuscript during its development.

## Author Contributions

Maggie MacPherson - concept, symposium organizer, symposium participant, writing & revising, editing full manuscript, topic editor (abstract, theory) and co-editor (temporal resolution of data types), figures, tables.

Kevin R. Burgio - concept, overall project support, symposium participant, writing & revising, conceptualizing and constructing figures, editing full manuscript, topic co-editor (methods & data types for spatial data)

Matthew G. DeSaix – symposium participant, writing & revising, topic co-editor (temporal resolution of data types)

Ben Freeman - symposium participant, writing, publication approach/ideas, topic editor (discussion)

Vitek Jirinec - symposium participant, writing & revising, topic co-editor (logistical methods and data types for occurrence data), figures.

John Herbert - symposium participant, writing, topic co-editor (methods & data types for spatial data)

Julia Shonfield - symposium participant, writing & revising, topic editor (introduction)

David Slager – symposium participant, writing & revising, topic editor (literature cited), topic co-editor (logistical methods and data types for occurrence data)

Rachael Herman - writing, topic co-editor (logistical methods and data types for occurrence data)

Charles van Rees – writing & revising, topic editor (analytical methods), figures.

Jill Jankowski – concept, symposium participant, writing & revising, editing full manuscript

## DATA ACCESSIBILITY STATEMENT

The R code for building Figure 1 is available at the authors github page [https://github.com/mmacphe/Global_Change_Biomes]. Data layer files were obtained from public online repositories.

## APPENDIX 1

Predicting future distributions from current relationships with explanatory variables is often done using freely available future climate projections (e.g., various “business as usual”, “middle of the road”, and “worst case scenario” models downloaded from https://www.worldclim.org), whereas hindcasting distributions is often done using paleoclimate models downloaded from the Intercomparison Project Phase II at https://www.pmip2.lsce.ipsl.fr. Here, we elaborate on the challenge of matching occurrence data with temperature and rainfall explanatory variable data sources because these are the two most widely used and generally important environmental factors in species distribution modeling (Bradie and Leung, 2017). The finest spatial and temporal resolution of environmental data is typically from weather station data from the nearest station to the research site (e.g., from https://www.wunderground.com/history) or by collecting weather data alongside occurrence data using data loggers available from various manufacturers (e.g., Meter, Truebner). The ERA5-Land dataset provides fine scale (hourly) temperature and precipitation data, but at a 9 km spatial resolution from January 1950 to present (https://cds.climate.copernicus.eu/cdsapp#!/dataset/reanalysis-era5-land?tab=overview). National Centers for Environmental Prediction (NCEP) provides global temperature and precipitation data (four-times daily) data at a 0.25 x 0.25 degree grid resolution from 1948-present (https://psl.noaa.gov/data/gridded/data.ncep.reanalysis.surface.html). WorldClim is a commonly used source of environmental data, and the current version (2.1) provides global monthly values of min and max temperature, and precipitation averaged from 1970-2000 (Fick and Hijmans, 2017). Historical monthly min and max temperature, and total precipitation data used for hindcasting are averaged by decade from 1960-2018 (Fick and Hijmans, 2017; Harris et al., 2014). BioClim data are 19 quarterly estimates derived from WorldClim monthly temperature and rainfall values that are thought to be more biologically meaningful than raw temperature or rainfall data (Busby, 1991). Despite being powerful tools for assessing macroclimate associations, caution should be used when using the most accessible climate products to inform conservation decisions by assessing the probability of future range locations with empirical validations that climate products (e.g., Bioclim variable 1 - annual mean temperature, Bioclim variable 5 - max temperature of the warmest month) are ecologically relevant to site occupancy.

